# Inflammatory Signatures of Microglia in the Mouse Model of Corticosterone-Induced Depression

**DOI:** 10.1101/2024.08.21.608928

**Authors:** Yanxiang Zhao, Yingying Huang, Ying Cao, Jing Yang

**Author notes:** Authors Contributed Equally (Yanxiang Zhao, Yingying Huang).

## Abstract

Microglia-mediated inflammation has been recognized as a key feature of major depressive disorder. Although hypercortisolemia commonly occurs in depressed patients and can be predictive of treatment response, how chronic exposure to this stress hormone influences microglia is incompletely characterized. Here, we exploited a standard mouse model of depressive-like behaviors induced by peripheral administration of corticosterone. Microglia in the prefrontal cortex of mice were profiled by bulk RNA sequencing, which exhibited the up-regulation of inflammatory markers. In addition, single-cell RNA sequencing identified distinct molecular patterns of microglial responses. Moreover, we revealed the elevation of Pu.1 and Irf8, the two central transcription factors governing microglia-mediated inflammation, in the prefrontal cortex and hippocampus of corticosterone-treated mice, which was similarly observed in the single-nucleus RNA sequencing dataset of depressed patients’ microglia. These results have established inflammatory signatures of microglia in the mouse model recapitulating hypercortisolemia-related depression, providing new insights into diagnostic and therapeutic strategies.

## Introduction

Major depressive disorder (MDD) is a psychiatric condition that severely disrupts the normal activities and daily life of patients [1–3]. Although the exact cause of MDD is yet to be fully understood, studies have documented inflammation as a key feature of MDD patients. In particular, plasma levels of inflammatory markers, e.g., C-reactive protein [4, 5] and interleukin (Il)-6 [6, 7], are elevated in a significant proportion of depressed patients. Accordingly, antidepressant treatments can effectively decrease such inflammatory markers while ameliorating MDD symptoms [8–10].

As the central type of residential immune cells in the brain, microglia are broadly involved in neurodevelopment and neurological diseases. For instance, microglia participate in the engulfment of apoptotic neurons and the pruning of synaptic connections during brain development, which represent vital steps in the precise establishment and maintenance of neurocircuits throughout adulthood [11–13]. Also, microglia can respond to diverse neurological conditions, e.g., neural damage and neurodegeneration. Microglia have long been known for their capacity to clear cellular debris left from neuronal or axonal death caused by traumatic neural injuries, which is indispensable for eliciting neuroinflammation and promoting tissue repair [14–17]. In addition, microglia are responsible for the phagocytic removal of protein aggregates such as Aβ plagues, which helps delay the onset and progression of Alzheimer’s disease [18–21]. On the other hand, microglia responding to neurodegenerative insults may release various inflammatory factors and aberrantly engulf synaptic structures, leading to collateral destruction of neurocircuits and exacerbation of disease conditions [22–24].

In contrast to those well-documented roles, microglial responses to psychiatric disorders have only begun to be recognized. In particular, accumulating evidence has indicated that microglia can participate in inflammatory events during depressive disorders. Microglial activation and proliferation are post-mortem observed in the brain regions involved in cognition and mood control, e.g., the prefrontal cortex and the hippocampus, of MDD patients [25–27]. In addition, similar microglial responses occur in those brain regions in the mouse models of depressive-like behaviors. Of importance, such microglia-mediated inflammation may alter the normal signals of neurotransmitters, e.g., dopamine, and neurocircuits involved in cognitive functions, thus contributing to the severity of depressive-like behaviors [28–30]. Indeed, it has been postulated that antidepressants such as ketamine may exert their therapeutic effects at least partially by modulating microglial inflammatory responses [31–33].

It has long been reported that depressed patients often have higher plasma levels of the stress hormone cortisol compared to healthy individuals [34–37]. This phenomenon of hypercortisolemia may be a consequence of chronic dysregulation of the hypothalamic-pituitary-adrenal (HPA) axis. While antidepressant medications can normalize hypercortisolemia in depressed patients, higher cortisol levels are predictive of more severe or treatment-resistant conditions [38–40]. However, recent research has implicated the complex causal relationship between cortisol and MDD [41–43]. In particular, hypercortisolemia may impede vital neurobiological processes in the brain, which can precede or even contribute to the development of depression. Notably, repeated peripheral administration of corticosterone in mice can reliably induce depressive-like behaviors [44, 45], e.g., increased immobility in the forced swim and tail suspension tests, inhibition of eating in a novel environment, and reduced exploration in the open field test. This corticosterone-induced mouse model also disrupts circadian rhythm [46], which similarly occurs in MDD patients with sleep disturbances [47, 48]. Despite those research advances, how chronic exposure to this stress hormone may influence microglial inflammatory responses remains incompletely characterized.

## Results

We exploited the standard mouse model recapitulating hypercortisolemia-related depression through the daily administration of corticosterone for three weeks. Microglia were then isolated by fluorescence-activated cell sorting (FACS) from the prefrontal cortex, a central brain region involved in depressive-like behaviors, of vehicle- or corticosterone-treated mice for transcriptomic analyses. As the entry point, two replicate preparations for each condition were subjected to bulk RNA sequencing (RNA-seq). Approximately 500 differentially expressed genes (DEGs) could be identified between the vehicle- *vs.* corticosterone-treated microglia (Fig. 1A and Table S1). Of importance, the expression of classic inflammatory markers, e.g., *Tnf*, *Il1a*, and *Il1b*, were among 173 up-regulated DEGs (Fig. 1B and Table S1). Also, several chemokines for immune cell recruitment, e.g., *Ccl2* and *Ccl3*, increased in the corticosterone-treated microglia (Fig. 1B). In addition, the signature genes of microglial activation, e.g., *Cd68* and *Apoe*, were evidently identified among up-regulated DEGs (Fig. 1B). On the other hand, the classic markers of resting “homeostatic” microglia, e.g., *P2ry12*, *Fcrls*, and *Sall1*, significantly decreased in the corticosterone-treated condition (Fig. 1B). These results implicated microglia-mediated inflammatory responses in this mouse model of depressive-like behaviors.

**Figure 1.**
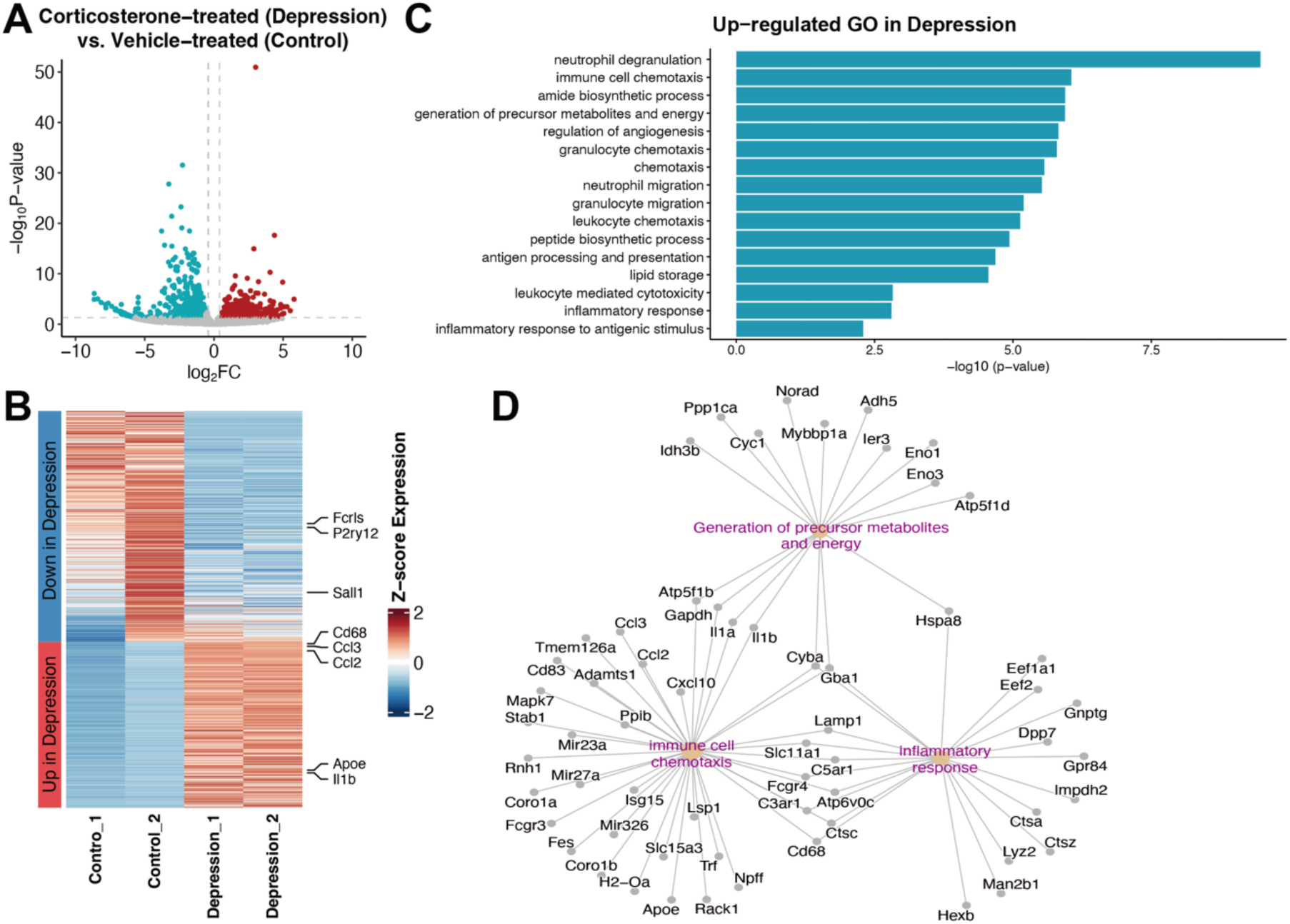
Inflammatory responses of microglia in the mouse model of corticosterone-induced depression. Microglia were FACS-sorted from the prefrontal cortex of vehicle-treated (control condition) or corticosterone-treated mice (depression condition) and subjected to bulk RNA-seq. **(A)** Volcano plot of differentially expressed genes (DEGs) identified in the corticosterone-treated microglia. **(B)** Heatmap plot of DEGs in the two replicate preparations of vehicle-treated or corticosterone-treated microglia. **(C)** GO enrichment and **(D)** GO network analysis of DEGs in the corticosterone-treated microglia.

We further compared the transcriptomics of the vehicle- *vs.* corticosterone-treated microglia by the Gene Ontology (GO) enrichment (Fig. 1C). In support of microglia-mediated inflammation, immune pathways were highly enriched in the corticosterone-treated condition. Also, pathways of key metabolic processes, e.g., protein synthesis and lipid storage, became up-regulated. Moreover, the GO network of those enriched processes revealed a collection of specific genes, e.g., *Il1b*, *Gapdh*, *Gba1*, and *Slc11a1*, that orchestrated such microglial responses (Fig. 1D).

For a more in-depth characterization of microglial responses in this mouse model of hypercortisolemia-related depression, we pursued the single-cell RNA sequencing (scRNA-seq) of microglia from the prefrontal cortex of vehicle- or corticosterone-treated mice. After quality control, a total of 21,818 cells (7,563 from vehicle-treated and 14,255 from corticosterone-treated) were obtained for detailed analyses. Six microglial clusters could be defined in the pooled scRNA-seq datasets (Fig. 2A and Fig. 2B), with the corticosterone-treated condition showing an apparent increase in clusters #3 and #4 (Fig. 2C and 2D). Notably, those microglial clusters exhibited significant inflammatory features. For instance, the inflammatory markers *Tnf* and *Il1b* both increased in the clusters #0 and #1 of corticosterone-treated microglia, and the *Il1b* up-regulation additionally occurred in the clusters #2, #3, and #4 (Fig. 2E), in line with the above observation with bulk RNA-seq data. Also, the expression of chemokines such as *Ccl2* became elevated in the microglial clusters #0, #2, #3, and #4 of depressive-like mice (Fig. 2E). In addition, the signature genes of microglial activation, e.g., *Cd68* (clusters #0, #1, #2, and #4), *Cd86* (clusters #0 and #1), and *Apoe* (clusters #0, #1, #2, #3, #4, and #5), were more enriched in the specific clusters of corticosterone-treated condition (Fig. 2E). These results thus revealed the distinct responses of microglial subpopulations in this mouse model of depression.

**Figure 2.**
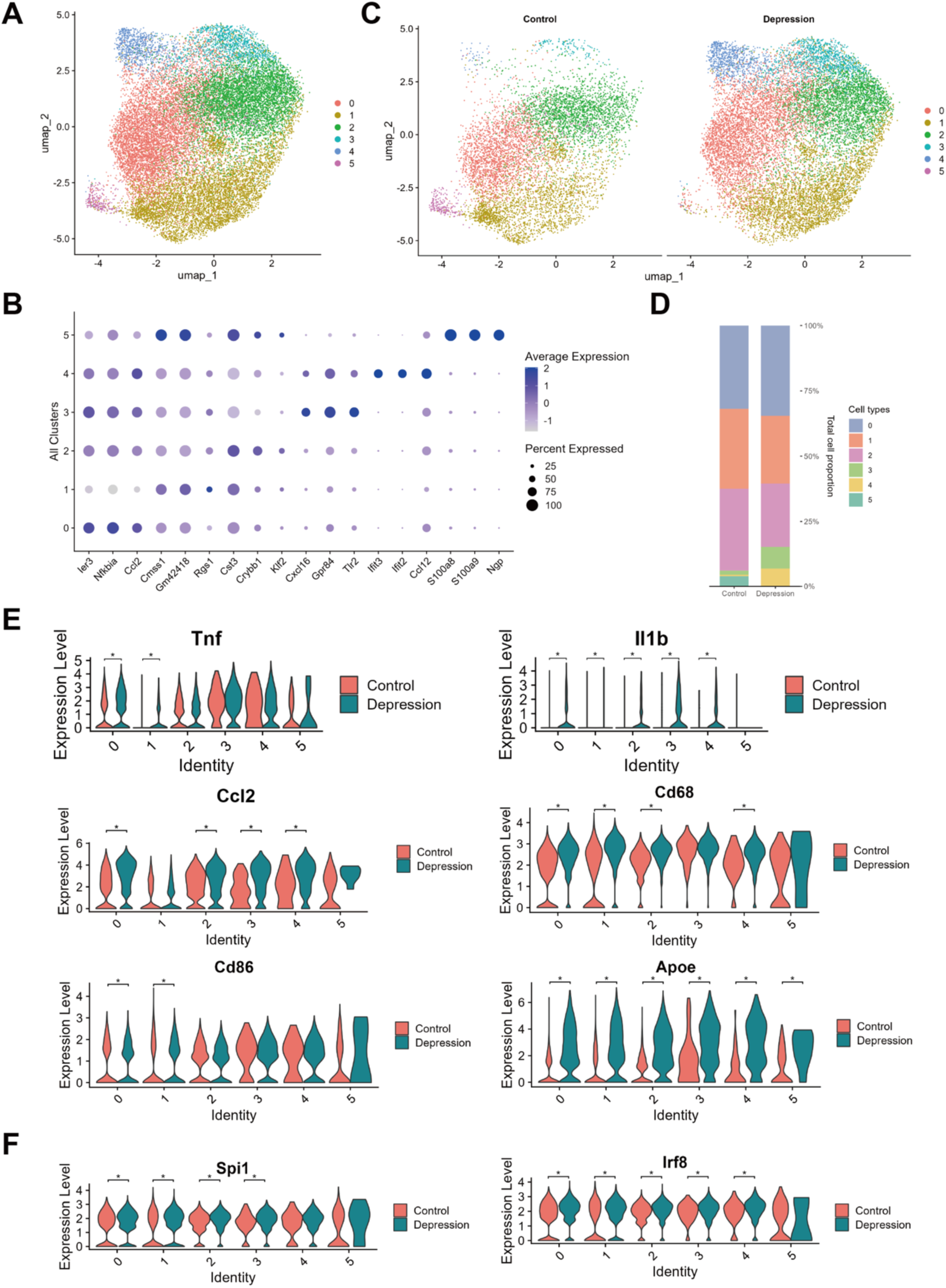
Inflammatory signatures of microglial subpopulations in the mouse model of corticosterone-induced depression. Microglia were FACS-sorted from the prefrontal cortex of vehicle-treated (control condition) or corticosterone-treated mice (depression condition) and subjected to scRNA-seq. **(A)** UMAP plot of microglial clusters (#0 ∼ #6) defined in the pooled scRNA-seq datasets. **(B)** Dot plot of the marker genes of defined microglial clusters. **(C)** UMAP plot of microglial clusters defined in the vehicle-treated or corticosterone-treated condition. **(D)** Percentage of each microglial cluster in the vehicle- *vs* corticosterone-treated condition. **(E)** Violin plots of inflammatory signature genes in the microglial clusters of vehicle- *vs* corticosterone-treated condition. **(F)** Violin plots of *Pu.1/Spi1* or *Irf8* expression in the microglial clusters of vehicle- *vs* corticosterone-treated condition. * *p* < 0.05 (Student’s *t*-test).

Previous studies by colleagues and us have established that the transcription factors Pu.1 (also known as Spi1) and Irf8 are central for microglial responses to diverse pathophysiological insults, e.g., traumatic neural injuries or neurodegenerative diseases [49–53]. In particular, these two transcription factors cooperatively instruct a repertoire of microglial function-related genes [52]. We observed in the bulk RNA-seq data that the corticosterone-treated microglia had increased levels of *Pu.1/Spi1* and *Irf8* (Table S1). Also, this up-regulation of *Pu.1/Spi1* (clusters #0, #1, #2, and #3) and *Irf8* (clusters #0, #1, #2, #3, and #4) could be detected in the majority of corticosterone-treated microglia by scRNA-seq (Fig. 2F).

We sought to verify the microglial up-regulation of Pu.1 and Irf8 in this mouse model of hypercortisolemia-related depression. Brain tissues of the vehicle- or corticosterone-treated mice were subjected to immunofluorescence staining of these two transcription factors. Indeed, expression levels of Pu.1 in Iba1^+^ microglia in the prefrontal cortex of corticosterone-treated mice became significantly elevated compared to the vehicle-treated condition (Fig. 3A and Fig. 3B). A similar increase of Pu.1 expression was simultaneously observed in the hippocampus (Fig. 3C and Fig. 3D), another key brain region implicated in depressive-like behaviors. In parallel, immunofluorescence staining showed the up-regulation of microglial expression of Irf8 in the prefrontal cortex (Fig. 3E and Fig. 3F) and the hippocampus (Fig. 3G and Fig. 3H) of corticosterone-treated mice. These results supported the involvement of Pu.1 and Irf8 in microglia-mediated inflammation in this mouse model of depression.

**Figure 3.**
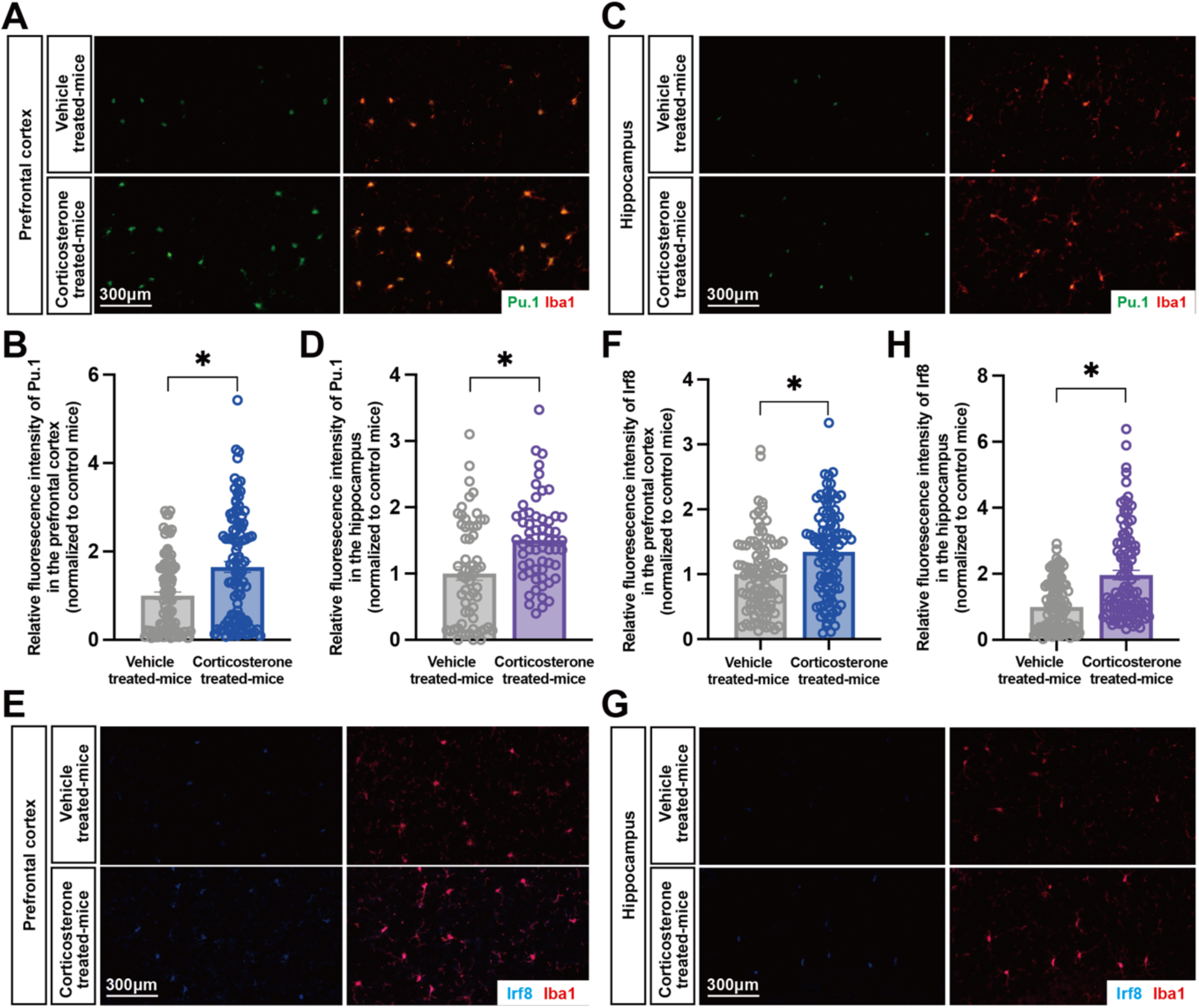
Microglial up-regulation of Pu.1 and Irf8 in the mouse model of corticosterone-induced depression. Brain tissues of vehicle-treated (control condition) or corticosterone-treated mice (depression condition) were subjected to immunofluorescence staining. **(A to D)** Co-staining of Pu.1 and Iba1 in mouse brain tissues. Representative images of the prefrontal cortex **(A)** or the hippocampus **(C)** were shown. Microglial intensities of Pu.1 immunofluorescence in the prefrontal cortex **(B)** or the hippocampus **(D)** were quantified. * *p* < 0.05 (Student’s *t*-test). **(E to H)** Co-staining of Irf8 and Iba1 in mouse brain tissues. Representative images of the prefrontal cortex **(E)** or the hippocampus **(G)** were shown. Microglial intensities of Irf8 immunofluorescence in the prefrontal cortex **(F)** or the hippocampus **(H)** were quantified. * *p* < 0.05 (Student’s *t*-test).

We explored the disease relevance of our transcriptomic analyses of microglia in the mouse model, taking advantage of the publicly available databases PsyGeNET [54] and DisGeNET [55, 56]. In PsyGeNET, 39 DEGs in the microglia of corticosterone-treated mice showed linkage to mental illness in the literature. Among a total of 135 associations, the highest number (61 associations) was identified for several categories of depressive disorders (Fig. 4A). In further support of this notion, a collection of gene signatures, including several inflammatory cytokines or chemokines, of corticosterone-treated microglia exhibited strong associations with MDD-related terms in DisGeNET (Fig. 4B).

**Figure 4.**
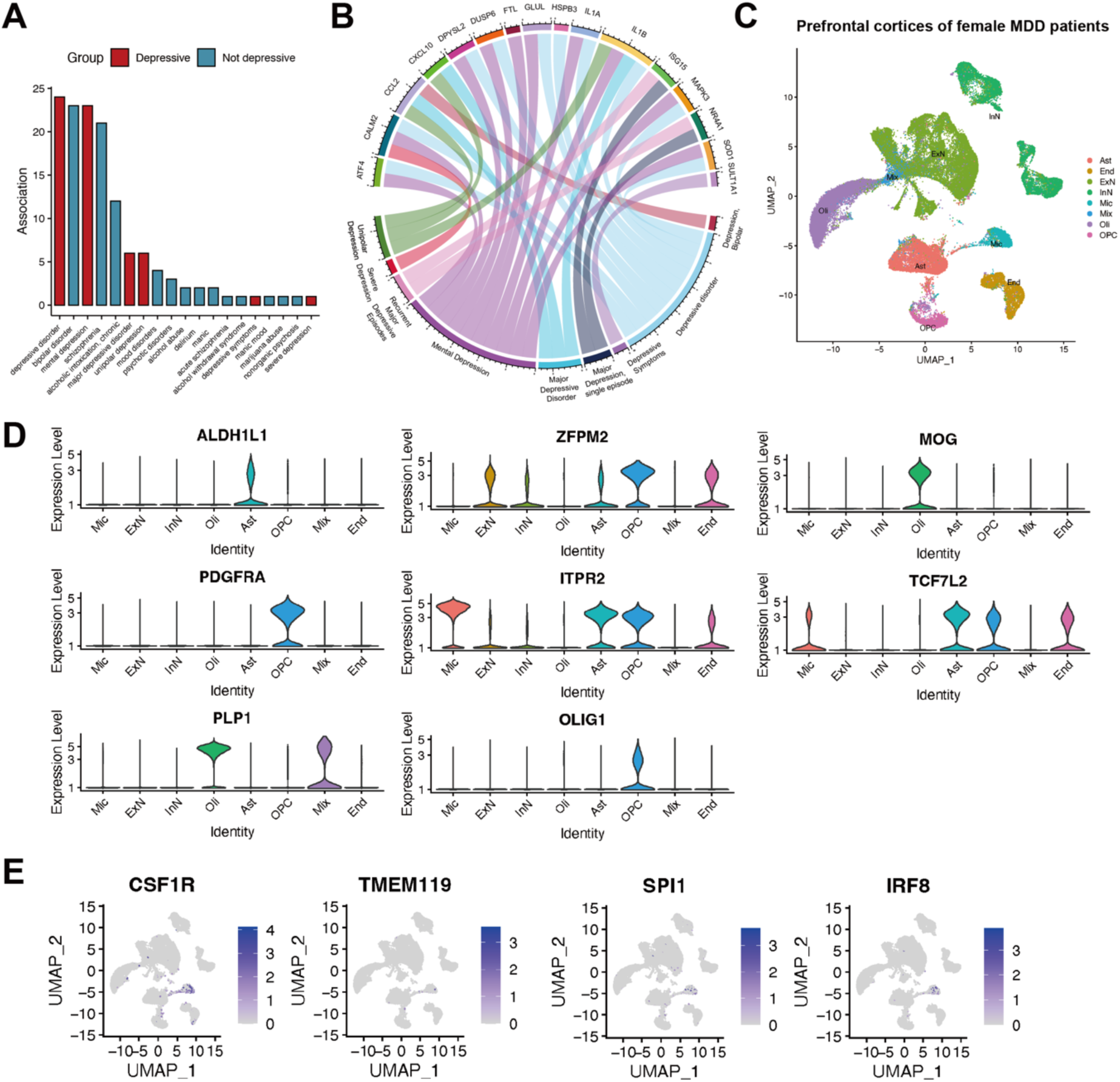
Potential involvement of PU.1 and IRF8 in microglial responses of MDD patients. **(A)** Bar graph of the associations of DEGs in corticosterone-treated microglia with mental illness in PsyGeNET. **(B)** Circos plot of the associations of DEGs in corticosterone-treated microglia with MDD- related terms in DisGeNET. **(C to D)** The published snRNA-seq dataset (GSE213982) of total cells in the prefrontal cortices of female MDD patients. **(C)** UMAP plot of neuronal and non-neuronal cell types defined in the dataset. **(D)** Violin plots of specific marker genes in the defined cell types of prefrontal cortices. **(E)** Feature plots of *CSF1R*, *TMEM119*, *PU.1/SPI1*, and *IRF8* in the defined cell types of prefrontal cortices of MDD patients.

We went on to examine the potential involvement of Pu.1 and Irf8 in microglia-mediated inflammation of MDD patients, looking into the single-nucleus RNA sequencing (snRNA-seq) dataset (GSE213982) of total cells in the prefrontal cortices of 20 female MDD patients [57]. Besides neuronal populations such as excitatory and inhibitory neurons, several non-neuronal cell types could be clearly defined, e.g., oligodendrocytes, astrocytes, and microglia (Fig. 4C and Fig. 4D). As a verification of the snRNA-Seq identification of microglia, the specific markers *CSF1R* and *TMEM119* were exclusively present in this cell cluster (Fig. 4E). Of importance, we observed that *PU.1* and *IRF8* showed enriched expression in the microglial population (Fig. 4E). These results have implicated the involvement of these two transcription factors in the microglia-mediated inflammation of depressed patients.

## Discussion

Recent studies have suggested that microglia-mediated inflammation represents a key feature of depression and potentially contributes to the disease’s onset and progression both in human patients and in mouse models [25–30]. Notably, while hypercortisolemia is commonly observed in depressed patients and may be predictive of their treatment responses, microglial inflammatory responses to chronic exposure to this stress hormone were incompletely determined. By exploiting the mouse model of recapitulating hypercortisolemia-related depression, this current work of transcriptomic analyses has revealed the distinct inflammatory signatures of microglia in the prefrontal cortex. Our identification of inflammatory markers and microglial subpopulations may provide a valuable reference for more precise diagnosis and stratification of MDD in future research. In addition, research attention is warranted to explore whether the condition of hypercortisolemia-related depression may share any common feature of microglial responses to other psychiatric disorders, e.g., autism spectrum disorder, schizophrenia, or drug addiction. Answers to this exciting question hold the promise of our better knowledge of the pathophysiological mechanisms of those debilitating human diseases.

Our colleagues and we have demonstrated the central role of transcription factors Pu.1 and IRF8 in microglial responses to various neurological conditions [49–53]. As a significant expansion from the previous findings, this study has revealed for the first time that the Pu.1-Irf8 signal may participate in microglia-mediated inflammation in both the mouse model of depression and MDD patients. How chronic cortisol exposure engages the Pu.1-Irf8 transcriptional module in microglia is currently unclear. A tempting possibility is that the cortisol-activated glucocorticoid receptor might directly target the gene expression of *Pu.1* or *Irf8*, which can be determined by future genomic analyses, e.g., chromatin immunoprecipitation sequencing (ChIP-seq) or cleavage under targets & tagmentation (CUT&Tag). In addition, whether the Pu.1-Irf8 signal similarly instructs microglial responses in other psychiatric diseases awaits clinical investigations. Furthermore, this Pu.1-Irf8 transcriptional module may open up a therapeutic target of microglia-mediated inflammation against depressive disorders, which calls for more translational research.

In sum, this study has reported the inflammatory signatures of microglia in the mouse model recapitulating hypercortisolemia-related depression, offering insights into the development of diagnostic and therapeutic strategies against this dreadful psychiatric disorder.

## Acknowledgments

This work has been supported by the National Natural Science Foundation of China (#32125017 and #32150008 to J.Y.), the National Key Research and Development Program of China (#2019YFA0802003 to J.Y.), the Beijing Natural Science Foundation (#7232086 to J.Y; #7234378 to Y.C.), and the China Postdoctoral Science Foundation (#2022M710219 to Y.C.). Y.C. was also supported by the Postdoctoral Fellowship of the Center for Life Sciences at Peking University. Additional funds to J.Y.’s research group have been from the State Key Laboratory of Membrane Biology at Peking University and the Center for Life Sciences at Peking University. The authors declare no conflict of interest.

## Author Contributions

Y.Z. and J.Y. designed the project. Y.Z., Y.H., and Y.C. performed experiments and analyzed the results. Y.Z., Y.H., and J.Y. prepared the manuscript.

## Materials and Methods

Further information and requests for resources or reagents should be directed to and will be fulfilled by the corresponding author, Jing Yang.

### Mouse information

All the experimental procedures in mice were performed in compliance with the protocol approved by the Institutional Animal Care and Use Committee (IACUC) of Peking University. Mice were maintained on the 12-hr/12-hr light/dark cycle (light period 7:00 am ∼ 7:00 pm), with the chow diet and water available *ad libitum*. 8-week-old C57BL/6 female mice were purchased from Charles River International.

For the mouse model of hypercortisolemia-related depression, corticosterone (Sigma) was formulated in DMSO / Kolliphor-EL (Sigma) / 5% sucrose (v:v:v = 1:3:6) and daily administered via oral gavage to C57BL/6 female mice at 50mg/kg of body weight for 3 weeks.

### Fluorescence-activated cell sorting (FACS) of microglia

The prefrontal cortical tissues of mice were freshly dissected and minced into small pieces on ice. The tissues were digested in RPMI 1640 (Thermo Fisher Scientific) / 0.1mg/ml Liberase TL (Roche) / 20μg/ml DNase I (Sigma-Aldrich) / 10mM HEPES / 3% heat-inactivated fetal bovine serum (HI-FBS; Sigma-Aldrich) at 37°C for 15 min. The tissues were mashed through a 70-μm cell strainer and centrifuged at 500 *g* for 5 min. The cells were re-suspended in PBS / 2% HI-FBS / 10mM EDTA-Na (pH 8.0) and stained with PE/Cyanine7 anti-mouse CD45 (BioLegend, #103114, RRID:AB_312979) and FITC anti-mouse/human CD11b (BioLegend, #101206, RRID:AB_312788). FACS-stained cells were processed on the Beckman MoFlo Astrios EQ, and microglia were sorted as CD45^low^ CD11b^+^ by the common standard.

### RNA sequencing (RNA-seq)

FACS-sorted mouse microglia were subjected to single-end RNA-seq by the Beijing Genomics Institute. Gene expression levels were normalized as fragments per kilobase per million (FPKM). The bulk RNA-seq data was deposited to the Sequence Read Archive (https://www.ncbi.nlm.nih.gov/sra) with the accession number GSE274360.

### Single-cell RNA sequencing (scRNA-seq)

FACS-sorted microglia were subjected to scRNA-seq by Singleron. The scRNA-seq data was deposited to the Sequence Read Archive (https://www.ncbi.nlm.nih.gov/sra) with the accession numbers GSM8447877 (vehicle-treated) and GSM8447878 (corticosterone-treated).

Singleron dataset processing was performed by CeleScope (v1.13.0) (https://github.com/singleron-RD/CeleScope), with mm10 as the reference mouse genome. The scrubletR (v0.1.0) (https://github.com/Moonerss/scrubletR) pipeline was applied to each dataset for removing potential doublets (min_counts=2, min_cells=3, expected_doublet_rate=0.06, min_gene_variability_pctl=85, n_prin_comps=50, sim_doublet_ratio=2). The filtered cell-by-gene count matrix was loaded into Seurat (v5.0.1) (https://github.com/satijalab/seurat) for in-depth analyses. We filtered out the genes expressed in <5 cells, as well as the cells expressing <600 or >5000 genes, >10% mitochondrial reads, or >20% ribosomal genes. We loaded the filtered count matrix to the CreateSeuratObject function to create a Seurat object, followed by the log-normalization by the NormalizeData function. The top 2,000 variable genes were identified using the FindVariableFeatures function. PCA was performed using the RunPCA function, and batch effect correction was then conducted on the principal components with the Harmony function to integrate two datasets. Unsupervised clustering was performed by the functions FindNeighbors and FindClusters. UMAP plots were generated by the RunUMAP function to visualize cell clusters.

### Single-nucleus RNA sequencing (snRNA-Seq)

The published snRNA-Seq dataset (GSE213982) of total cells in the prefrontal cortices of 20 female MDD patients [57] were obtained from GEO (https://www.ncbi.nlm.nih.gov/geo/). For quality control, we applied filters for genes expressed in 10 or fewer cells, cells expressing fewer than 200 genes, mitochondrial reads exceeding 10%, or ribosomal genes for more than 20% of the expression. We further employed Solo in scvi-tools to remove doublets, using the top 2000 variant genes and default parameters. We then utilized single-cell Variational Inference (scVI) in scvi-tools to integrate the snRNA-Seq data of different patients, following the software instructions (https://docs.scvi-tools.org/en/stable/tutorials/index.html). SCANPY was used for clustering the integrated data, and the resulting H5ad files were converted into the Seurat objects using Sceasy (https://github.com/cellgeni/sceasy). Finally, we performed the analyses in Rstudio with R v4.2.25, utilizing Seurat v.4’s FindAllMarkers to determine and visualize the markers for each cluster. For the determination of cell types, we referred to PanglaoDB [58] and the cell identity information provided in the original datasets. The feature plots were generated using scCustomize (https://samuel-marsh.github.io/scCustomize/).

### Immunofluorescence staining

The mice of indicated conditions were perfused with PBS / 50μg/ml heparin followed by PBS / 1% PFA / 50μg/ml heparin. The brain tissues were harvested and post-fixed in PBS / 1% PFA at room temperature for 2hr and then cryopreserved in PBS / 30% sucrose at 4°C overnight for 10-μm cryosectioning. The sections were immunolabeled with the intended primary antibodies (final concentration of 1μg/ml) in PBS / 0.1% Tween-20 / 5% normal donkey serum. The primary antibodies used for mouse tissue sections were rabbit anti-Pu.1 (Cell Signaling Technology, Cat#2258, RRID:AB_10693421), rabbit anti-Irf8 (Cell Signaling Technology, Cat#98344, RRID:AB_3083757), and rat anti-Iba1 (Abcam, Cat# ab283346, RRID:AB_3065282). The tissue sections were further immunolabeled with the corresponding secondary antibodies Alexa Fluor 488-conjugated donkey anti-rat IgG (Thermo Fisher Scientific, #A21208, RRID:AB_2535794) or Alexa Fluor 568-conjugated donkey anti-rabbit IgG (Thermo Fisher Scientific, #A10042, RRID:AB_2534017).

The immunostained tissue sections were scanned by Axio Scan Z1. Microglia and their indicated features were quantified in ImageJ (https://imagej.net/ij).

### Statistical methods

Student’s *t*-tests (two-tailed unpaired) were performed using GraphPad Prism 9.5.0 (http://www.graphpad.com/scientific-software/prism). Statistical details of the experiments are included in figure legends.

